# Common neural mechanisms supporting time judgements in humans and monkeys

**DOI:** 10.1101/2024.04.25.591075

**Authors:** Julio Rodriguez-Larios, Elie Rassi, Germán Mendoza, Hugo Merchant, Saskia Haegens

## Abstract

There has been an increasing interest in identifying the biological underpinnings of human time perception, for which purpose research in non-human primates (NHP) is common. Although previous work, based on behaviour, suggests that similar mechanisms support time perception across species, the neural correlates of time estimation in humans and NHP have not been directly compared. In this study, we assess whether brain evoked responses during a time categorization task are similar across species. Specifically, we assess putative differences in post-interval evoked potentials as a function of perceived duration in human EEG (N = 24) and local field potential (LFP) and spike recordings in pre-supplementary motor area (pre-SMA) of one monkey. Event-related potentials (ERPs) differed significantly after the presentation of the temporal interval between “short” and “long” perceived durations in both species, even when the objective duration of the stimuli was the same. Interestingly, the polarity of the reported ERPs was reversed for incorrect trials (i.e., the ERP of a “long” stimulus looked like the ERP of a “short” stimulus when a time categorization error was made). Hence, our results show that post-interval potentials reflect the perceived (rather than the objective) duration of the presented time interval in both NHP and humans. In addition, firing rates in monkey’s pre-SMA also differed significantly between short and long perceived durations and were reversed in incorrect trials. Together, our results show that common neural mechanisms support time categorization in NHP and humans, thereby suggesting that NHP are a good model for investigating human time perception.

## Introduction

Time estimation in the range of hundreds of milliseconds is a crucial ability for many species, as it is necessary for a wide variety of behaviours including foraging and communication. The last decade has seen an increasing interest in the identification of neural underpinnings of motor and perceptual timing (Balasubramaniam et al., 2021; Tsao et al., 2022). Neurophysiological experiments have suggested the existence of a core timing network that includes the medial premotor areas (SMA and preSMA) and its (sub) cortical connections (Merchant et al., 2013). The current hypothesis is that the medial premotor cortex encodes both elapsed time and the temporal scaling of its neural population trajectories in state space (Gámez et al., 2019; Sohn et al., 2019). These dynamics are linked to the existence of neural sequences that form patterns of active neurons changing in rapid succession and that flexibly cover an interval depending on the timed duration (Crowe et al., 2014; Merchant et al., 2015).

Electroencephalography (EEG) is an ideal tool to investigate the neural correlates of time estimation in humans due to its high temporal resolution. For this purpose, previous studies have combined EEG with different interval timing tasks (Bueno & Cravo, 2021; Damsma et al., 2021; Duzcu, 2019; Lindbergh & Kieffaber, 2013; Ng et al., 2011; Özoğlu & Thomaschke, 2023; Pfeuty et al., 2005). Most of these tasks involve the presentation of visual or auditory stimuli to signal a to-be-timed interval, which has to be compared to a reference time interval or prototype. Early studies focused on ERPs during the presentation of the time interval (Macar & Vidal, 2003; Pfeuty et al., 2005). However, it has been recently shown that time estimation is better reflected in post-interval potentials (i.e., evoked responses emerging after the offset of the time interval; Kononowicz & Penney, 2016; Kononowicz & Rijn, 2014). In this way, it has been shown that post-interval potentials in fronto-central electrodes differ significantly depending on the perceived time duration (Damsma et al., 2021; Kononowicz & Rijn, 2014; Kruijne et al., 2021; Lindbergh & Kieffaber, 2013; Özoğlu & Thomaschke, 2023; Tarantino et al., 2010).

Although research in NHP has provided insights into the neural correlates of time estimation (Merchant et al., 2013), these have not yet been directly compared to electrophysiological findings in humans. This is primarily because research in NHP is often focused on spiking activity (Leon & Shadlen, 2003; Mendoza et al., 2018), which cannot be recorded non-invasively in healthy human participants. Although no previous study has investigated post-interval potentials during time estimation tasks in monkeys, they are expected to be qualitatively similar to those found in humans. Indeed, the behaviour of monkeys and humans on time estimation tasks shows similar psychometric properties, which suggests a common neural substrate (Mendez et al., 2011; Zarco et al., 2009). Moreover, evoked potentials have been reported in monkeys in other cognitive tasks, showing similar dynamics to the ones observed in humans (Godlove et al., 2011; Peissig et al., 2007).

Here, we assess whether the neurophysiological signatures of time perception are similar in humans and NHP. For this purpose, we analysed EEG data from 24 humans and extracellular recordings in pre-SMA from one monkey while they performed a temporal interval categorization task. In this task, participants had to decide whether a visually presented temporal interval had a shorter or longer duration than a previously learned prototype. Based on prior work, we hypothesized that post-interval evoked potentials would significantly differ between “short” and “long” perceived durations in both humans and NHP, and that this would be accompanied by changes in firing rates in the latter.

## Methods

### Participants

*Humans*. 27 healthy adult subjects (12 males) participated in the experiment. The mean age was 25.6 years old (SD = 4.2). Participants reported normal or corrected-to-normal vision and no history of neurologic or psychiatric diagnosis. Informed consent procedure and study design were approved by the Institutional Review Board (IRB) of the New York State Psychiatric Institute (protocol #8001). Participants were compensated for their participation (at 25 USD per hour). Three participants were excluded from the analysis due to technical problems during data acquisition.

Monkeys. One male Rhesus monkey (Macaca mulatta; 5.5 kg) was tested. All experimental procedures were approved by the National University of Mexico Institutional Animal Care and Use Committee and conformed to the principles outlined in the Guide for Care and Use of Laboratory Animals (NIH, publication number 85–23, revised 1985).

### Stimuli and task

*Humans*. Participants performed a temporal interval categorization task (see Figure 1A). In this task, participants had to categorize a visual stimulus based on its duration on screen. Each trial started with the presentation of a fixation cross for 2 seconds. Then, the to-be-categorized stimulus (a circle around the fixation cross) was shown for a specific time interval. After another 2 seconds delay, participants had to report whether the presented stimulus was “short” or “long” by pressing the right or left arrow key on a computer keyboard. In order to avoid motor preparation, response mappings (i.e., left vs. right arrow key) were randomly changed on a trial-by-trial basis. Feedback was presented at the end of each trial via a colored fixation cross (green for correct and red for incorrect responses). The task involved three stimulus sets (T1, T2 and T3) with different interval durations (see Figure 1B). This design allowed usto compare “short” and “long” decisions for stimuli with the same objective duration (see two black boxes in **Figure 1B**). The different sets were presented in a blocked design, with order randomised per participant. Each block had a learning phase of ten trials in which participants would learn the meaning of “short” and “long” durations for that set (only the shortest and the longest intervals of each block were presented). This allowed participants to implicitly learn the category boundary. A total of 336 trials (112 per block) were performed, with the experiment lasting for approximately 1 hour.

**Figure 1.**
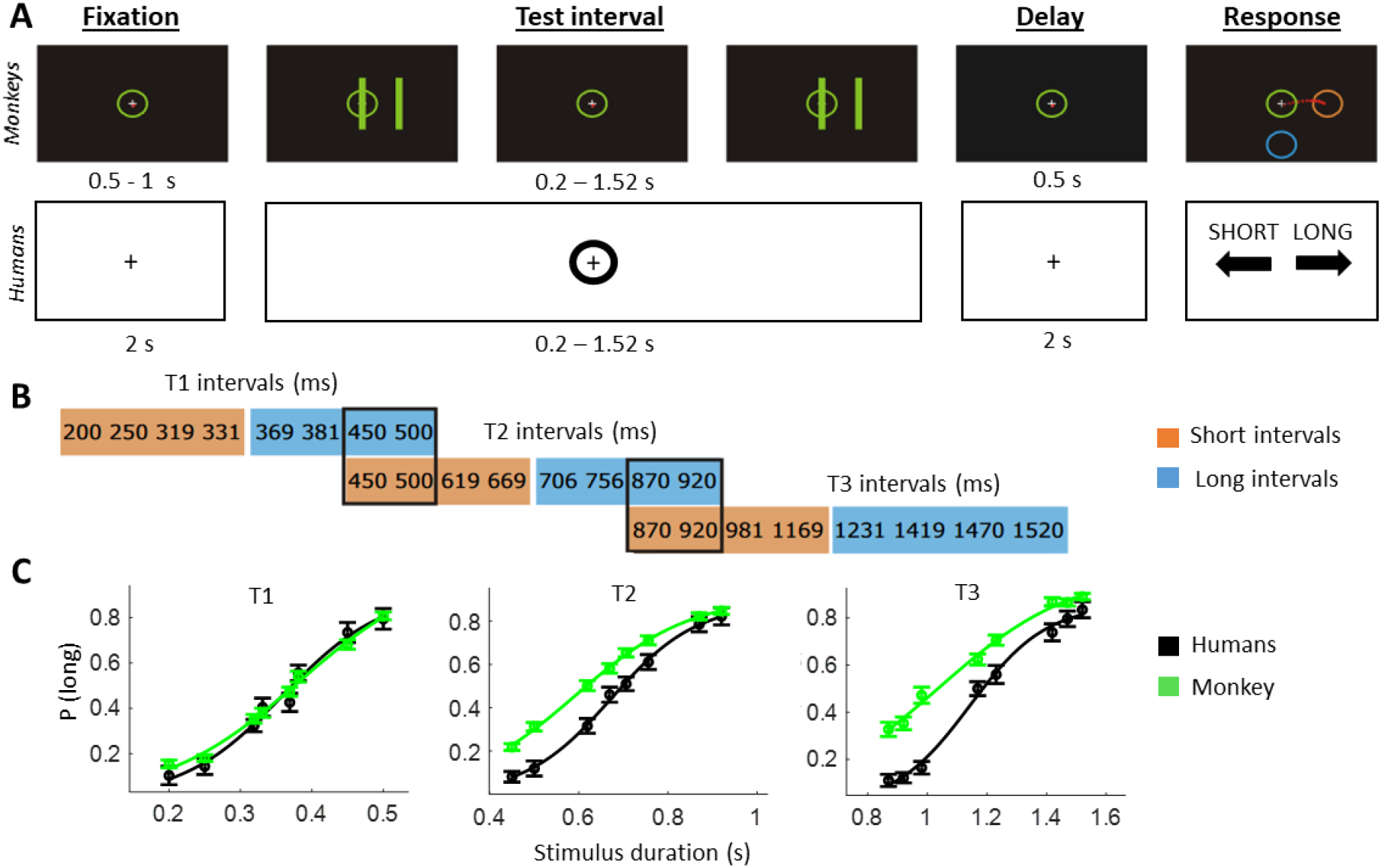
Temporal categorization task. A) Schematic of the time-interval categorization tasks adopted for monkeys (first row) and humans (second row). Subjects had to indicate whether the test interval was of “long” or “short” duration. Response mapping was randomized per trial and shown after the decision delay. B) Interval durations for each of the three sets. Note that certain intervals could be “short” in one set but “long” in another (marked with black outline). C) Psychometric curves (probability of answering “long”) for humans (black) and one monkey (green), per stimulus set. Error bars represent standard error across subjects for humans and across sessions for the monkey.

*Monkeys*. Task details have been reported previously (Mendoza et al., 2018). In short, monkeys were trained to categorize the temporal interval between two visual stimuli as either “short” or “long,” according to previously learned prototypes. First, a circle containing a fixation point was shown in the center of the screen. Then, the animal started the trial by staring at the fixation point and by placing the cursor inside the central circle. After a variable waiting period (500–1000 ms), two parallel bars separated by constant distance appeared for 50 ms, disappeared for a particular test interval, and reappeared in the same position. The first and second stimulus presentations indicated the beginning and the end of the test interval, respectively. After a fixed delay (500 ms) two response targets (orange and blue circles) were presented. Both response targets could occupy one of eight possible locations on the periphery of the screen. The monkeys were trained to move the cursor from the central circle to the orange target if the test interval was short or to the blue target if it was long. The monkey received a juice reward immediately after each correct response. The task involved three stimulus set s (T1, T2 and T3) with different interval durations (see Figure 1B), presented in separate trial blocks. Each block had an initial instruction phase of 24 trials in which only the shortest and the longest intervals of each block were presented. In these trials the color of the parallel bars matched the color of the correct response target (orange for the short interval and blue for the long interval). The following 96 trials constituted the test phase in which the color of the bars was green regardless of the stimulus category. A total of 199 sessions (each of them involving 96 experimental trials) were performed.

### Recordings

*Humans*. 96-electrode scalp EEG was collected using the BrainVision actiCAP system (Brain Products GmbH, Munich, Germany) with a sampling rate of 500 Hz. Electrodes were labeled according to the international 10-20 syste. The reference electrode during the recording was Cz. Amplification and digitalization of the EEG signal was done through an actiCHamp DC amplifier (Brain Products GmbH, Munich, Germany) linked to BrainVision Recorder software (version 2.1, Brain Products GmbH, Munich, Germany). Vertical (VEOG) and horizontal (HEOG) eye movements were recorded by placing additional bipolar electrodes above and below the left eye (VEOG) and next to the left and right eye (HEOG).

*Monkeys*. Recording chambers (8-mm inner diameter) were implanted over the left pre-SMA and dorsolateral prefrontal cortex (dlPFC) during aseptic surgery under Sevoflurane (1–2%) gas anesthesia. Chamber positions were determined on the basis of structural MRI. Titanium posts for head restraining were implanted on the skull. Broad spectrum antibiotics (Enrofloxacin, 5 mg/kg/day, i.m.) and analgesics (Ketorolac 0.75 mg/kg/6 h or Tramadol 50–100 mg/4–6h,i.m.) were administered for 3 days after surgery. The extracellular activity of neurons in pre-SMA was recorded with quartz-insulated tungsten microelectrodes (1–3MΩ) mounted in multielectrode manipulators (Eckhorn System, Thomas Recording, GMbH, Giessen, Germany). All neurons were recorded regardless of their activity during the task, and the recording site changed from session to session. Spike waveform data were sorted online employing window discriminators (Blackrock Microsystems LLC, Salt Lake City, UT, USA). LFP data were simultaneously recorded from both pre-SMA and dlPFC using a 250-Hz low-pass filter and stored at 1,000 Hz for offline analysis. The titanium posts of the head-restraining implant were used for grounding.

### Pre-processing of electrophysiological data

*Humans*. Pre-processing was performed in MATLAB R2021a using custom scripts and functions from EEGLAB (Delorme & Makeig, 2004) and Fieldtrip (Oostenveld et al., 2011) toolboxes. Data were first resampled to 250 Hz and filtered between 0.5 and 30 Hz. Noisy electrodes were automatically detected (EEGLAB function clean_channels) and interpolated. EEG data were re-referenced to the common average and independent component analysis (runica algorithm) was performed. An automatic component rejection algorithm (IClabel) was employed to discard components associated with muscle activity, eye movements, heart activity or channel noise (threshold = 0.8)(Pion-Tonachini et al., 2019). In addition, components with an absolute correlation with HEOG, VEOG or ECG channels higher than 0.8 were discarded. Furthermore, Artifact Subspace Reconstruction (ASR) was employed to correct for abrupt noise with a cut-off value of 20 SD(Chang et al., 2019) ERPs were obtained by averaging trials within subjects, condition and electrodes. In order to reduce the dimensionality of the data, Principal Component Analysis (PCA) was used to compute spatial filters that explained most of the variance of the EEG data(Guarnieri et al., 2020; Zanotelli et al., 2010). We concatenated ERPs across subjects to compute common spatial filters for all subjects (i.e., Group PCA analysis) (Dien, 2012).

*Monkeys*. All LFP pre-processing was done with Fieldtrip and custom MATLAB R2019a code. Only data from pre-SMA was used based on prior work (Kononowicz & Penney, 2016). Epochs were visually inspected and excessively noisy channels and trials were rejected (around 10 % of data)(Rassi et al., 2023). Data were filtered between 0.5 and 30 Hz. ERPs were obtained by averaging trials within pre-SMA electrodes, sessions and conditions.

### Statistical analysis

For the behavioural analysis, we calculated psychometric curves per subject (in humans) or per session (in monkeys). For this purpose, the probability of categorizing each interval of the corresponding set of stimuli as “long” was fitted with a logistic function, which was defined as:

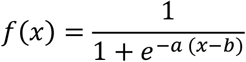

The parameter *a* represents the steepness of the slope and *b* is the sigmoid midpoint. The slope was extracted per subject (in humans) or per session (in monkeys) to be statistically compared between sets and species. These comparisons were done using ANOVAs and t-tests (paired and independent) as implemented in MATLAB.

For the electrophysiological data, a cluster-based permutation test (Maris & Oostenveld, 2007) was used to assess condition-related statistical differences in ERPs. This test controls for the type I error rate arising from multiple comparisons by using a non-parametric Montecarlo randomization and taking into account the dependency of the data. First, cluster-level statistics were estimated in the original data and in 1,000 permutations of the data. Cluster-level statistics are defined as the sum of t-values with the same sign across adjacent time points that are above a specified threshold (i.e., 97.5th quantile of a t-distribution). Then, the cluster-level statistics from the original data were evaluated using the reference distribution obtained by taking the maximum cluster t-value of each permutation. Cluster-corrected p-values are defined as the proportion of random partitions whose cluster-level test statistic exceeded the one obtained in the observed data. The significance level for the cluster permutation test was set to 0.025 (corresponding to a false alarm rate of 0.05 in a two-sided test). A paired-samples t-test was chosen as the first-level statistic to compare experimental conditions in humans and monkeys.

## Results

### Behaviour

The psychometric curves of both the human participants and the monkey followed a typical sigmoid shape showing that the probability of categorizing a particular interval as ‘‘long’’ increased as a function of the interval duration (see **Figure 1C**).

The slope of the psychometric curves, which reflects sensitivity, differed significantly between sets in both the monkey (F (2,134) = 51.93; p < 0.001) and humans (F (2,69) = 5.54; p = 0.006). In line with previous literature (Mendez et al., 2011), post-hoc t-tests showed that the slope of the psychometric curve became flatter in blocks with longer stimulus duration, thereby reflecting decreased sensitivity with longer intervals in both species. Specifically, post-hoc t-tests showed that in humans the slope of the psychometric curve was significantly flatter in set T2 relative to set T1 (t(23) = 3.30; p = 0.003) and in set T3 relative to set T1 (t(23) = 3.27; p = 0.003), but not in set T3 relative to set T2 (t(23) = 0.93; p = 0.35). In the monkey, the slope of the psychometric curve was significantly flatter in set T3 relative to both set T1 (t(86) = 8.47; p < 0.001) and set T2 (t(83) = 3.43; p < 0.001), and in set T2 relative to set T1 (t(99) = 7.07; p < 0.001). When directly comparing the slope of the psychometric curve between species, we found that humans had greater sensitivity (i.e., steeper slope) than monkeys in set T1 (t(74) = 2.64; p = 0.01), set T2 (t(71) = 3.67; p < 0.001) and set T3 (t(59) = 2.39; p = 0.020).

In sum, these behavioural results indicate that both humans and the monkey successfully categorized time intervals and that the psychometric properties of behavioural responses were similar across species.

### Post-interval evoked responses in humans

We first sought to replicate and extend previous findings showing differences between “long” and “short” temporal decisions in post-interval ERPs of human EEG. In order to reduce the dimensionality of the data, we performed a group PCA. We selected the first three components for further analysis since they cumulatively explained over 70% of the variance (40.3 %, 24.9 % and 11.7 %, respectively). Only the first two components showed significant differences between “short” and “long” decisions (see below).

The first principal component (**Figure 2A**) showed a more pronounced positive potential around 300 ms for correct “short” decisions (*t*_*cluster*_ = -113.89; *p*_*cluster*_ = 0.002) and a more pronounced negative potential around 600 ms for correct “long” decisions (*t*_*cluster*_ = -100.59; *p*_*cluster*_ = 0.002; **Figure 2B**). The same pattern of results was observed when selecting correct trials with the same objective duration (i.e., matched for physical stimulus properties but belonging to a different category), although in this case, only the difference around 300 ms remained significant (*t*_*cluster*_ = -95.64; *p*_*cluster*_ = 0.003; **Figure 2C**). No significant differences were found for incorrect trials (**Figure 2D**).

**Figure 2.**
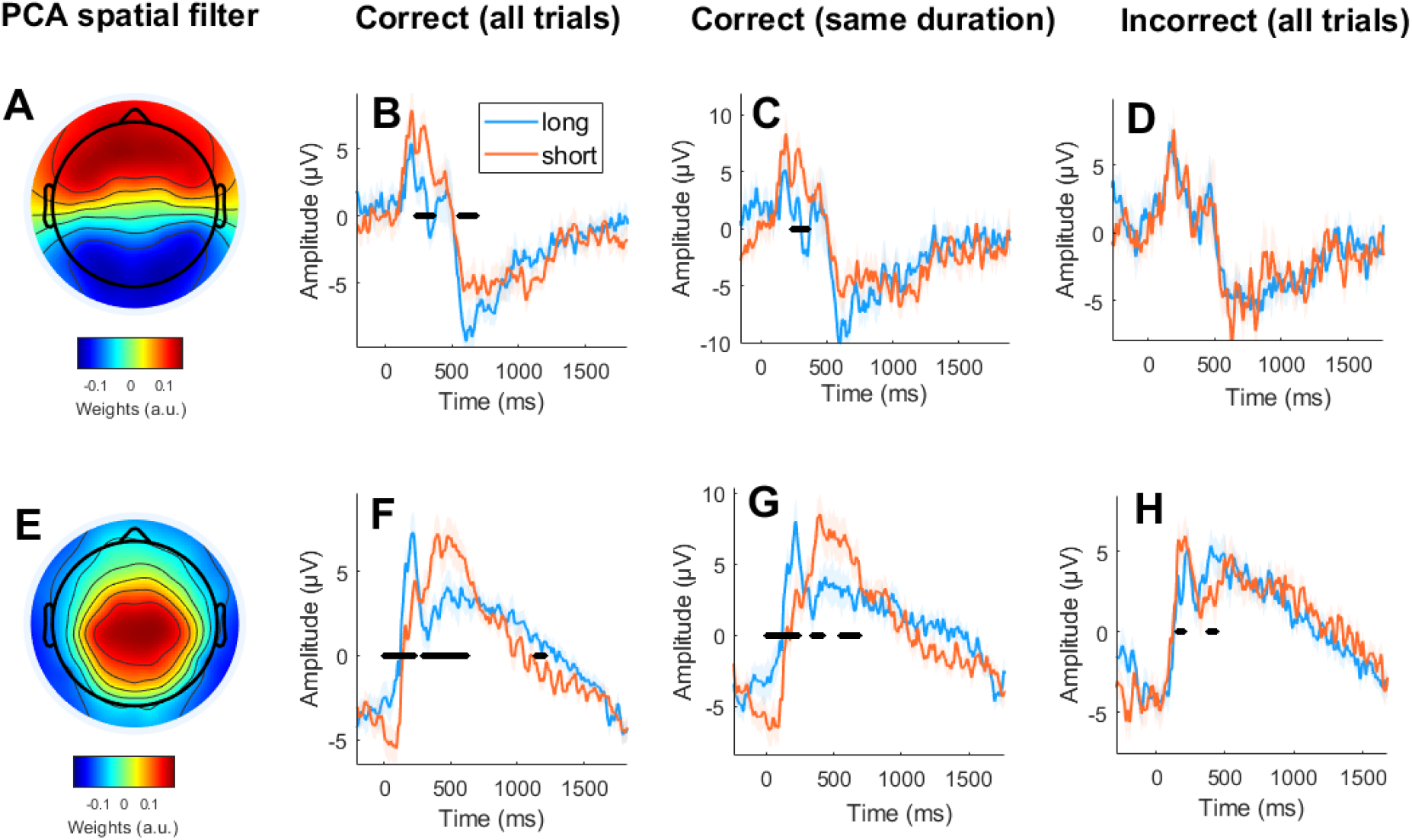
Post-interval evoked potentials in humans. Each row depicts the ERPs after a Group PCA-derived spatial filter is applied. (A) Topography of the spatial filters for the first principal component. (B) ERPs for “long” and “short” responses for correct trials. (C) ERPs for “long” and “short” responses for correct trials with the same objective (but differently perceived) duration. (D) ERPs for “long” and “short” responses for correct trials. (E-H) same as A-D for second component. Statistical significance at p<0.025 is marked with a black line, and 0 on the x-axis represents the offset of the stimulus whose duration was to be evaluated.

The second principal component (**Figure 2E**) showed a more pronounced positive potential around 200 ms for “long” decisions (*t*_*cluster*_ = 231.63; *p*_*cluster*_ < 0.001), a more pronounced positive potential around 400 ms for “short” decisions (*t*_*cluster*_ = -283.94; *p*_*cluster*_ < 0.001), and a more pronounced negative potential around 1000 ms for “short” decisions (*t*_*cluster*_ = 64.89; *p*_*cluster*_ = 0.024; **Figure 2F**). The same pattern of results was observed when selecting correct trials with the same objective duration (*t*_*cluster*_ = 189.78, *p*_*cluster*_ = 0.002; *t*_*cluster*_ = -99.44, *p*_*cluster*_ = 0.003; *t*_*cluster*_ = -71,03, *p*_*cluster*_ = 0.009; Figure 2G). Crucially, this pattern of results was reversed for incorrect trials for the ERPs around 200 ms and 400 ms (*t*_*cluster*_ = 49.91, *p*_*cluster*_ = 0.009 and *t*_*cluster*_ = -45.96, *p*_*cluster*_ = 0.018, respectively; Figure 2H).

Together, these results show that post-interval potentials reflect the perceived duration of the stimuli as i) ERPs for “short” and “long” decisions differed significantly even if the objective duration of the stimulus was the same, and ii) some of these ERP effects were reversed for incorrect trials.

### Post-interval responses in human and non-human primates

In order to compare post-interval potential differences between “long” and “short” temporal decisions across species, we selected a group of frontal electrodes in humans that overlaps spatially with the recording locations in the monkey (**Figure 3AE**).

**Figure 3.**
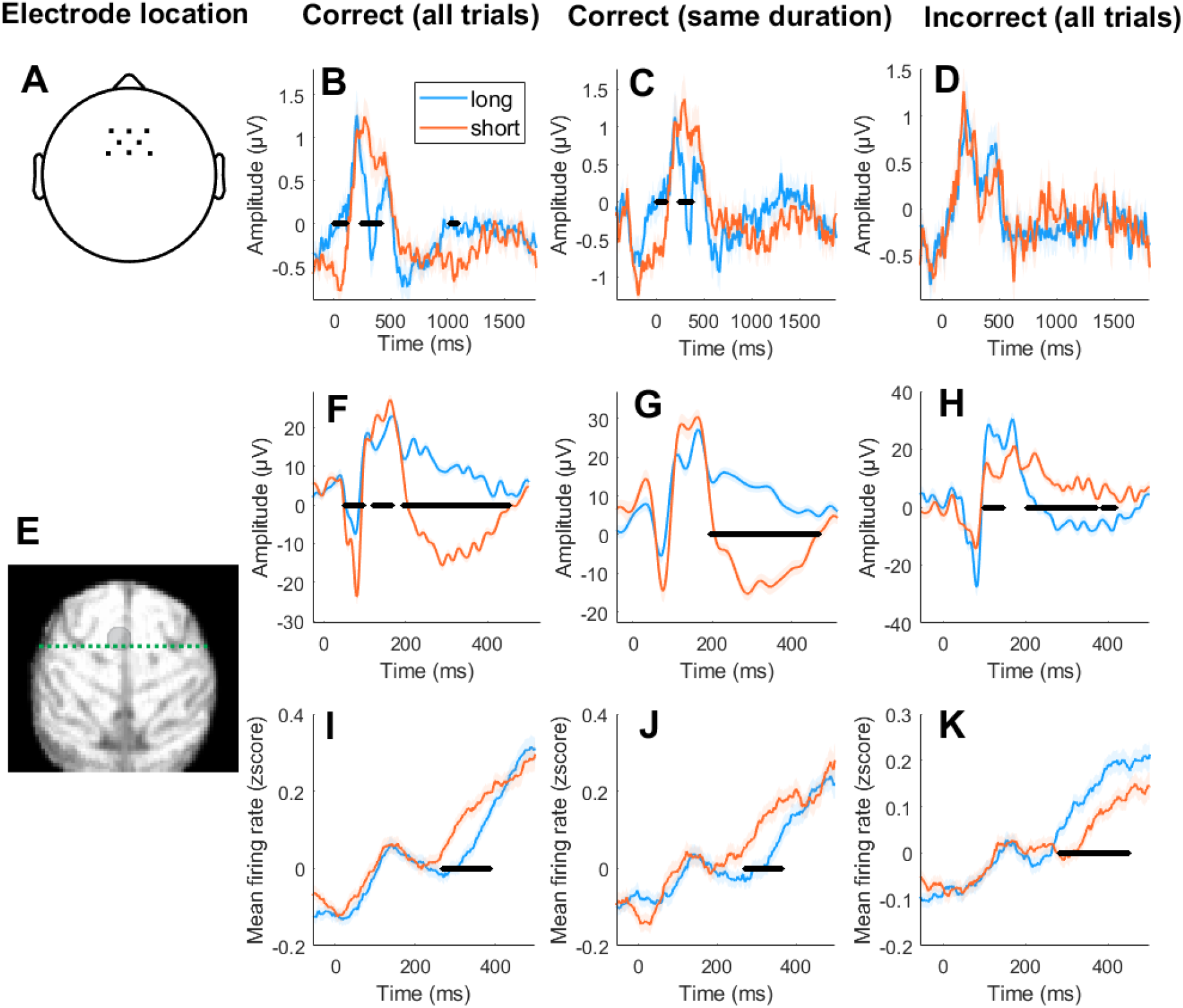
Fronto-central post-interval evoked responses in human and non-human primates. (A) Topography showing the location of electrode selection in humans. (B) ERPs for “short” (orange) and “long” responses (blue) in humans for correct trials. (C) ERPs for “short” (orange) and “long” responses (blue) in humans for correct trials with the same objective (but differently perceived) duration. D) ERPs for “short” (orange) and “long” responses (blue) in humans for incorrect trials. (E) Analysed recording sites in the monkey (pre-SMA). (F-H) Same as B-D but for the monkey. (I-K) Same as B-D but depicting monkey’s firing rate instead of ERPs. Statistical significance at p<0.025 is marked with a black line in each subplot, and 0 on the x-axis represents the offset of the stimulus whose duration was to be evaluated.

In humans, correct “short” decisions were associated with a more pronounced negative potential in the first 100 ms (*t*_*cluster*_ = 123.64; *p*_*cluster*_ = 0.002), a more pronounced positive potential around 200 ms (*t*_*cluster*_ = -166.69; *p*_*cluster*_ < 0.001) and a more pronounced negative potential around 1000ms (*t*_*cluster*_ = 56.42; *p*_*cluster*_ = 0.021; **Figure 3B**). A qualitatively similar pattern of results was observed when selecting trials with the same objective (but differently perceived) duration (*t*_*cluster*_ = 68.53; *p*_*cluster*_ = 0.016; *t*_*cluster*_ = -108.58; *p*_*cluster*_ = 0.003; **Figure 3C**). No significant differences were identified for incorrect trials (**Figure 3D**).

Similarly, correct “short” decisions in the monkey were associated with a more pronounced negative potential in the first 100 ms (*t*_*cluster*_ =13475; *p*_*cluster*_ <0.0001), a more pronounced positive potential around 200 ms (*t*_*cluster*_ = -71.18; *p*_*cluster*_ = 0.004) and a more pronounced negative potential around 300 ms (*t*_*cluster*_ =120.28; *p*_*cluster*_ = 0.004; **Figure 3F**). A qualitatively similar pattern of results was observed when selecting trials with the same objective (but differently perceived) duration (*t*_*cluster*_ = 1147; *p*_*cluster*_ < 0.001; **Figure 3G**). Moreover, this pattern of results was significantly reversed for incorrect trials (*t*_*cluster*_ = -71.96; *p*_*cluster*_ < 0.001; *t*_*cluster*_ = -333.35 ; *p*_*cluster*_ < 0.001; **Figure 3H**). In addition, relative firing rate in the monkey was more pronounced for correct “short” decisions around 300 ms for all trials (*t*_*cluster*_ = -197.68; *p*_*cluster*_ = 0.002; **Figure 3I**) as well as for trials matched for duration (*t*_*cluster*_ = -139.48; *p*_*cluster*_ = 0.004; **Figure 3J**), and this pattern of results was reversed for incorrect trials (*t*_*cluster*_ = 252.63; *p*_*cluster*_ < 0.001; **Figure 3K**).

Together, these results show that: i) post-interval potentials in monkey pre-SMA and human frontocentral EEG are qualitatively similar and reflect perceived time durations, and ii) some of these differences are also mirrored in firing rates modulations in the monkey.

## Discussion

In this study, we assessed whether monkeys and humans share the same neural mechanisms for time estimation. For this purpose, we analysed the electrophysiological signals of 24 humans (EEG) and one monkey (extracellular recordings in pre-SMA) while they categorised a temporal interval as either “short” or “long” based on previously learned prototypes. Our results show that evoked potentials after the presentation of the time interval differed significantly between “short” and “long” decisions in both humans and monkey. Crucially, we show that these differences reflect the perceived (and not the objective) duration of the time intervals because: i) the same difference in post-interval potentials was evident when stimuli had the same objective (but differently perceived) duration, and ii) the reversed pattern of results was observed for incorrect trials. In addition, some of the differences in post-interval potentials were accompanied by significant changes in monkey’s firing rates.

Previous literature has shown significant differences in post-interval ERPs as a function of the perceived duration in human EEG. Specifically, when the presented stimulus is perceived as longer than the reference, a more pronounced positive potential around 200 ms (P200) has been found, while for shorter stimuli a more pronounced Late Positive Potential (LPP) and P300 have been found (Damsma et al., 2021; Kononowicz & Rijn, 2014; Kruijne et al., 2021; Lindbergh & Kieffaber, 2013; Özoğlu & Thomaschke, 2023; Tarantino et al., 2010). These potentials (P200, P300 and LPP) are thought to reflect comparison and decision processes (Kononowicz & Rijn, 2014; Lindbergh & Kieffaber, 2013). Our results replicate these findings in humans and show that these potentials are encompassed in two different components (P300 in the first PCA and P200/LPP in the second PCA), which suggests different neural generators (Dien, 2012). In addition to previously reported post-interval potentials, we found significant differences between “short” and “long” decisions in a slow negative potential (occurring after 500 ms). This slow negative potential peaked around 600 ms for “long” perceived durations and around 1000 ms for “short” perceived durations (**Figure 2B**). Based on previous literature (Bosch et al., 2001), we interpret these slow frontocentral negative potentials as a neural mechanism supporting working-memory retention after the decision is made.

It has been proposed that the differences in post-interval potentials as a function of perceived duration observed in humans reflect differences in the timing of cognitive processes supporting time estimation, rather than a timing mechanism in itself (Kononowicz & Penney, 2016; Kononowicz & Rijn, 2014; Lindbergh & Kieffaber, 2013). It can be argued that when the interval is longer than the category boundary, subjects can make a decision before the interval offset. For stimuli shorter than the category boundary, participants are only able to decide after the interval offset (Mendoza et al., 2018). This interpretation is supported by differences in the ERPs reported here. Specifically, comparison/decision processes could be reflected in the more pronounced P200 for long decisions and in the later P300/LPP for short decisions (**Figure 2BF**). Memory retention of the decision could be reflected in the negative slow potential peaking at 600 ms for long decisions and at 1000 ms for short decisions.

In order to compare human and monkey post-interval neural responses, we selected a cluster of frontocentral electrodes in humans that overlapped with the location of the recordings in the monkey (around pre-SMA). Strikingly, the neural dynamics observed in both species were highly similar, showing significant differences between short and long perceived durations with the same polarity. In addition, the changes in the later post-interval potential in the monkey were mirrored in the firing rate, which suggests differential excitation levels in pre-SMA for short and long perceived durations. Since we show shared neural substrate of time categorization between humans and monkeys, our findings support the idea that research on the monkey brain can help elucidate the neural mechanisms supporting human time estimation, and, by extension, its related deficits in clinical populations (Merchant et al., 2008).

The main limitations of the current work are related to differences between species in terms of experimental design and recording methods used. First, the temporal categorization task was not exactly the same in the human and monkey experiments. The tasks differed in both the presentation of the temporal interval (empty interval in monkey vs. filled interval in human) and the duration of the post-interval delay (2 s in human and 0.5 s in monkey). Recordings in humans were done with scalp EEG, while recordings in monkeys were with intracortical electrodes. These factors may have affected the shape of the ERPs and should be controlled for in future research. To avoid possible confounders, future studies should: i) include EEG recordings in NHP, and ii) make sure that both species perform exactly the same task.

In conclusion, this study extends previous findings regarding post-interval evoked potentials in the context of time estimation in humans and shows that similar neural mechanisms are present in monkeys. Therefore, our results further support the idea that the monkey brain is a good model to investigate the neural mechanisms underlying human time perception.

## Author contributions

J.R. designed and performed the experiment, analysed the data and wrote the paper. E.R. helped to analyse the data. G.M. performed the experiment and helped analyse the data. H.M. designed and performed the experiment and edited the paper. S.H. supervised the project and edited the paper.

